# On the phenology of protists: Recurrent patterns reveal seasonal variation of protistan (Rhizaria: Cercozoa, Endomyxa) communities in tree canopies

**DOI:** 10.1101/2021.02.15.431229

**Authors:** Susanne Walden, Robin-Tobias Jauss, Kai Feng, Anna Maria Fiore-Donno, Kenneth Dumack, Stefan Schaffer, Ronny Wolf, Martin Schlegel, Michael Bonkowski

## Abstract

Tree canopies are colonized by billions of highly specialized microorganisms that are well adapted to the extreme microclimatic conditions, caused by diurnal fluctuations and seasonal changes. In this study we investigated seasonality patterns of protists in tree canopies of a temperate floodplain forest via high-throughput sequencing with group-specific primers for the phyla Cercozoa and Endomyxa. We observed consistent seasonality and identified divergent spring and autumn taxa. Tree crowns were characterized by a dominance of bacterivores and omnivores, while eukaryvores gained a distinctly larger share in litter and soil communities on the ground. Seasonality was largest among communities detected on the foliar surface. Higher variance within alpha diversity of foliar communities in spring indicated greater heterogeneity during community assembly. However, communities underwent distinct changes during the aging of leaves in autumn, reflecting recurring phenological changes during microbial colonization of leaves. Surprisingly, endomyxan root pathogens appeared to be exceptionally abundant across tree canopies during autumn season, demonstrating a potential role of the canopy surface as an important reservoir for wind-dispersed propagules. Overall, about 80% of detected OTUs could not be assigned to known species – representing only a fraction of dozens of microeukaryotic taxa whose canopy inhabitants are waiting to be discovered.

## INTRODUCTION

### Tree canopies – an ephemeral environment for microbes

The forest canopy is defined as ‘the aggregate of all tree crowns in a stand of vegetation, which is the combination of all foliage, twigs, fine branches, epiphytes as well as the air in a forest’ (Parker, Lowman and Nadkarni 1995). With an estimated area exceeding 100 million km^2^ globally, the foliar surface forms the largest biological surface on earth (Morris and Kinkel 2002; Peñuelas and Terradas 2014). Nevertheless, knowledge on microorganisms inhabiting the phyllosphere, i.e. the whole aerial region of plants dominated by leaves (Vorholt 2012), is far less advanced than that of below-ground counterparts. The phyllosphere is subject to extreme microclimatic dynamics due to rapid changes in abiotic stressors such as UV radiation, temperature, humidity and osmotic pressure during daily fluctuations that only specially adapted microorganisms can cope with (Baldocchi and Collineau 1994; Vorholt 2012; Manching, Balint-Kurti and Stapleton 2014; Stone, Weingarten and Jackson 2018). Considering that perennial deciduous plants produce and shed their leaves every year, the phyllosphere represents a highly ephemeral environment (Vorholt 2012; Mwajita *et al.* 2013). Thus, it can be presumed that microorganisms dwelling within this habitat opportunistically colonize, multiply and occupy newly formed niches after leaf emergence throughout the year.

### Seasonal variability – a major shaping agent of foliar bacterial communities

Former studies on foliar microecology observed bacteria to be by far the most abundant inhabitants, with on average 10^6^-10^7^ bacterial cells per cm^2^ of foliar surface (Lindow and Brandl 2003; Rastogi, Coaker and Leveau 2013). Investigations into the variation of microbial communities on leaves over multiple temporal and spatial scales provided detailed knowledge on the taxonomy and the ecology of bacterial leaf inhabitants (Thompson *et al.* 1993; Jacques, Kinkel and Morris 1995). Seasonal variability turned out to be a major driver of variation in these prokaryotic communities (Lauber *et al.* 2013). Another, but still neglected important factor shaping foliar bacterial communities are microbial predators, i.e. bacterivorous protists (Mueller and Mueller 1970; Bamforth 1973, 2007, 2010; Flues, Bass and Bonkowski 2017). Protistan predation has a profound influence on the structure and function of bacterial communities (Matz and Kjelleberg 2005; Krome *et al*. 2010; Jousset 2012; Amacker *et al*. 2020). Since these microbial eukaryotes comprise a vast array of functional traits in morphologies, locomotion and feeding modes (Fiore-Donno *et al.* 2019; Dumack *et al.* 2020), we presume that different protistan taxa play contrasting or complementary ecological roles within the heterogeneous habitat of forest canopy.

### On the seasonal variability of protists

In contrast to molecular surveys on seasonal changes in prokaryote diversity (Rastogi *et al.* 2012; Copeland *et al.* 2015; Agler *et al.* 2016), studies on community shifts of protists over time were commonly conducted in aquatic systems for dominant taxa (Rynearson, Newton and Armbrust 2006; Aguilera *et al.* 2007) or at higher taxonomical level (Tamigneaux *et al.* 1997; Araújo and Godinho 2008); studies on terrestrial protists often lack a temporal dimension. Consequently, analyses of seasonality in terrestrial protistan communities are a rarity and hitherto limited to a relatively small range of ecosystem types, dominated by soil habitats (Fiore-Donno *et al.* 2019; Fournier *et al.* 2020). Hence, the effect of a seasonal niche separation as a possible selective force which causes seasonal shifts of protistan communities dwelling on plant surfaces remains largely unexplored.

### Protists and their distribution mechanisms

Dispersal of unicellular organisms in terrestrial environments is facilitated by dormant stages, i.e. resting cysts or spores (Foissner 1987, 2006; Verni and Rosati 2011). These can be carried over large distances by wind (Wilkinson 2001), rain and fog (Finlay 2002), or animals and humans (Revill, Stewart and Schlichting 1967; Schlichting and Sides 1969; Perrigo, Romeralo and Baldauf 2012). Recent studies on protists with taxon-specific primers allow for the first time a thorough recovery of the existing species richness in a habitat and indeed suggest a largely ubiquitous distribution within the same terrestrial ecosystem (Fiore-Donno *et al.* 2018, 2019; Degrune *et al.* 2019; Jauss *et al.* 2020a). Considering the large surface area that trees extend into the atmosphere, the forest canopy may act as huge reservoir for airborne microorganisms, thus may be conducive to their further spread into the surrounding soils (Jauss *et al.* 2020b). Accordingly, it may be suggested that community assembly is driven largely by random dispersal, but because the canopy is subject to extreme environmental conditions where only adapted species will successfully replicate and survive, we expect specific patterns of beta diversity to dominate over random community assembly. Moreover, the question arises whether protistan communities undergo further seasonal changes, forced by changing abiotic conditions, or after the colonization of newly formed leaves and subsequent successions towards adapted species.

In this study, we investigated seasonal changes in protistan communities of different microhabitat compartments in the canopy region of three autochthonous tree species in a temperate floodplain forest. We further compared the canopy communities to those of the litter layer and mineral soil on the ground. Four samplings were conducted in two consecutive spring and autumn seasons, over a period of two years. We applied a MiSeq Illumina sequencing protocol using taxon-specific primers for the protistan phyla Cercozoa and Endomyxa (Rhizaria) (Fiore-Donno, Richter-Heitmann and Bonkowski 2020). Cercozoa are a highly diverse group representing many taxa and encompassing a variety of functional traits, and Endomyxa are of particular interest for comprising diverse plant parasites of economic importance (Neuhauser *et al.* 2014; Bass, Ward and Burki 2019, Dumack *et al.* 2020).

We hypothesized that **(I)** cercozoan and endomyxan communities differ in their seasonal composition in tree canopies. **(II)** Functional diversity of communities differs spatially and temporally between different microhabitats. **(III)** Despite the presumption that tree canopies act as a reservoir for wind-dispersed propagules, we expect specific patterns of beta diversity to dominate over randomness in communitiy assembly throughout all samplings.

## MATERIAL AND METHODS

### Sampling, DNA extraction and sequencing

Microhabitat samples were collected during spring and autumn within a period of two years: October 2017 and 2018, and May 2018 and 2019. The sampling took place in cooperation with the Leipzig Canopy Crane Facility in the floodplain forest in Leipzig, Germany (51.3657 N, 12.3094 E). All samples were obtained and processed as described in Jauss et al. (2020a). Briefly, seven different microhabitat compartments were sampled related to the canopy surface at 20-30m height: fresh leaves, deadwood, bark, arboreal soil and three cryptogamic epiphytes comprising lichen, and two moss species, *Hypnum* sp. and *Orthotrichum* sp. For comparison, two samples on the ground (leaf litter layer and mineral soil underneath up at to 10 cm depth) were sampled. All microhabitat samples were taken with four treatment replicates from three tree species *(Quercus robur, Tilia cordata* and *Fraxinus excelsior)* with three biological replicates each. DNA extraction was done according to the manufacturer’s protocol with the DNeasy PowerSoil kit (QIAGEN, Hilden, Germany). DNA concentration and quality were checked using a NanoDrop Spectrophotometer (NanoDrop Technologies, Wilmington, USA). Semi-nested PCRs with tagged group-specific primers (Fiore-Donno, Richter-Heitmann and Bonkowski 2020) and Illumina sequencing were performed as described in Jauss et al. (2020a), the used primer and barcode combinations are provided in Supplementary Table S1 and S2.

### Sequence processing

Sequence processing followed the pipeline described in Fiore-Donno, Richter-Heitmann and Bonkowski (2020). Briefly, paired reads were assembled using MOTHUR v.39.5 (Schloss *et al.* 2009) allowing no differences in the primer and the barcode sequences and no ambiguities. Next, assembled sequences smaller than 300bp and with an overlap less than 200bp were removed. The obtained sequences were checked for their quality and clustered into Operational Taxonomic Units (OTUs) using VSEARCH (Rognes *et al.* 2016) with abundance-based greedy clustering (agc) and a similarity threshold of 97%. Clusters represented by ≤0.005% of the total number of reads were removed to reduce amplification errors and sequencing noise (Fiore-Donno *et al.* 2018). Sequences were assigned with the PR2 database (Guillou *et al.* 2013) using BLAST+ (Camacho *et al.* 2009) with an e-value of 1^e-50^, keeping only the best hit. Cercozoan an endomyxan sequences were aligned with a template provided in Fiore-Donno et al. (2018). Finally, to detect chimeric sequences UCHIME (Edgar *et al.* 2011) was used as implemented in MOTHUR.

To explore the sequencing depth by sample metadata, the final OTU table was loaded into QIIME2 v2018.11 (Bolyen *et al.* 2019). To ensure sufficient sequencing depths for further analyses a threshold for a minimum number of sequences per sample was determined, which was set as high as possible: at least five samples per microhabitat and 15 samples per tree species (≤7525 reads sample^−1^).

### Functional traits

We classified the protistan OTUs according to their respective feeding modes into bacterivores, eukaryvores and omnivores (i.e. feeding on both bacteria and eukaryotes) as in Dumack et al. (2020). The phytomyxean parasites, due to their peculiar life cycle, were considered separately in each functional category. We assigned traits at the genus level (Supplementary Table S3).

### Statistical analyses

All statistical analyses were conducted in R v3.5.3 (R Core Team, 2019). Rarefaction curves were carried out with the iNEXT package (Hsieh, Ma and Chao 2015) to determine if a higher sequencing depth would have revealed more diversity. Alpha diversity indices were calculated for each microhabitat per sampling period using the *diversity* function in the vegan package (Oksanen *et al.* 2019). Analysis of season correlated OTU abundances was performed with the DESeq2 package (Love, Huber and Anders 2014) at the 1% significance level. To explore differences in community composition between the samples, the following beta diversity-based methods were conducted on relative abundances: Non-metric multidimensional scaling was performed on the Bray-Curtis dissimilarity matrix (functions *vegdist* and *metaMDS* in the vegan package); to show differences between fresh leaves communities of different sampling periods a principal coordinate analysis (PCoA, function *cmdscale* in the vegan package) was performed; to analyse the effects of environmental factors on the variance of the community composition, a redundancy analysis was carried out on the Hellinger-transformed table (function *rda* in the vegan package); to test if protistan OTUs and functional diversity differed across the sampled strata, microhabitats, tree species and seasons, a permutational multivariate analysis of variance (perMANOVA, function *adonis*) and, where appropriate, an analysis of variance (ANOVA, function *aov*) were conducted. The number of shared OTUs between different combinations of microhabitats was visualized using the UpSetR package (Lex *et al.* 2014; Gehlenborg 2019). Figures were plotted with the ggplot2 package (Wickham, 2016). Cercozoan and endomyxan diversity was illustrated using the Sankey diagram generator (http://sankeymatic.com/, 12 December 2020, date last accessed).

## RESULTS

### Sequencing results

We obtained 783 genuine cercozoan and endomyxan OTUs from 324 canopy and ground microhabitat samples representing on average 1.5 million filtered sequences per sampling period and 6 157 731 high quality sequence reads in total (Supplementary Table S4). However, 34 samples (ca. 10%) were removed because the yield was not sufficient (≤7525 reads sample^−1^). The remaining 290 samples yielded on average 20 657 reads sample^−1^ (min. 7633; max 57 404; SD 9520). The average number of OTUs was 780 ± 1, 781 ± 2 and 774 ± 1 per microhabitat, tree species and sampling period, respectively. In total 22% of the OTUs showed a sequence similarity of 97-100% to any known reference sequence (Figure 1 B). OTU001 occurred with exceptionally high read abundances in the canopy, being 18-fold higher than in the ground stratum (1 183 933 vs. 67 009 reads; ANOVA: F = 68.98, p < 0.001, Figure 1A). Whereby, OTU001 had 86.14% sequence similarity to a molecularly undetermined glissomonadid species (Figure 1 A; Supplementary Table S5).

**Figure 1:**
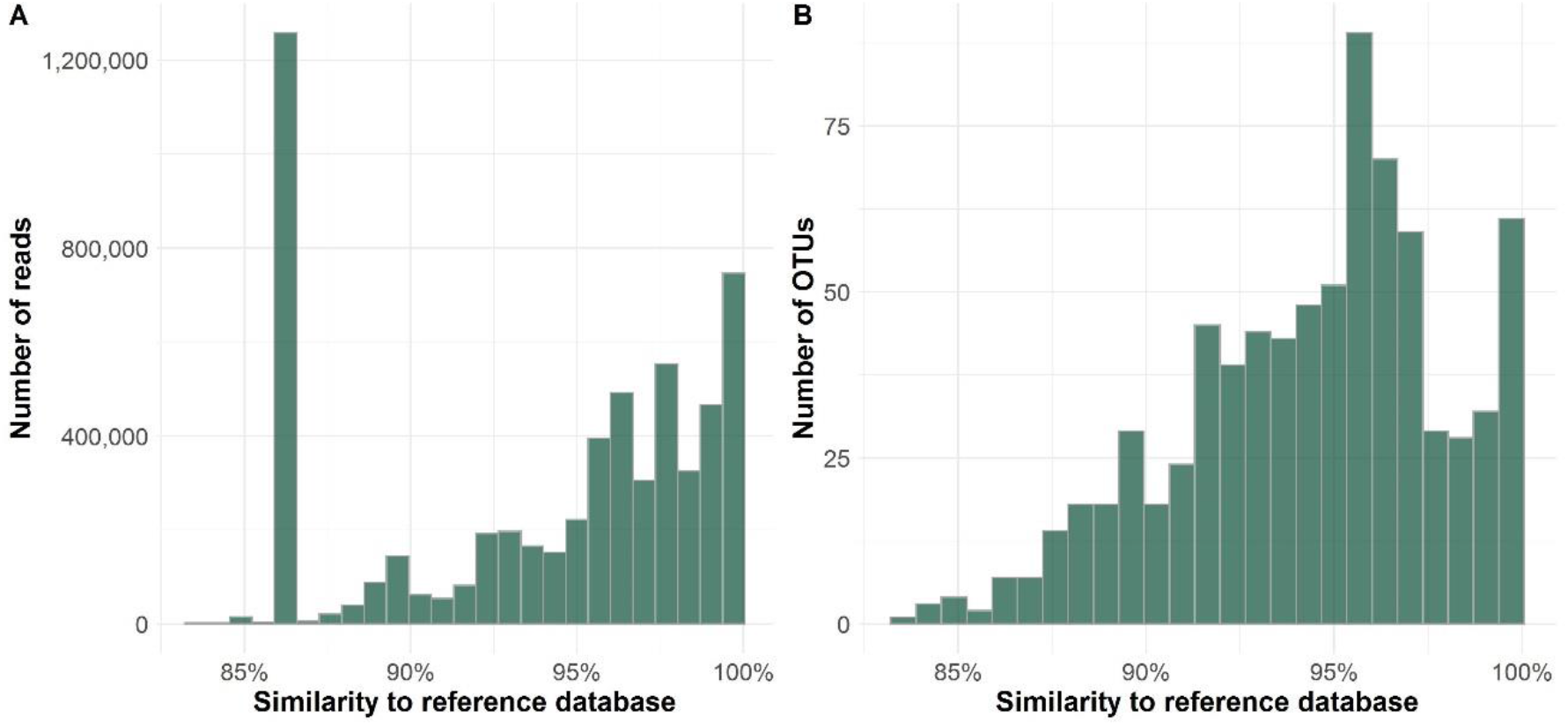
Similarity of protistan reads and OTUs to the reference database. Only 37% of all reads (A) and 22% of all OTUs (B) were ≥97% similar to sequences within the respective database. Read numbers of OTU001 (long bar in Figure 1A) exceed more than 1 million reads in tree canopies and was the most abundant OTU in every sampling period.

Sampling effort was sufficient for the majority of sampled microhabitats in both autumn samplings, where the total OTU richness was reached after only ca. 200 000 sequences. In spring samples, however, rarefaction curves for several microhabitats did not reach a plateau, especially for the samples of fresh leaves (Supplementary Figure S1), suggesting that we underestimated the OTU richness in this habitat. A database with OTU abundances, taxonomic assignment and functional traits is provided (Supplementary Table S3).

### Seasonal variation and spatial structuring

Investigation into seasonality patterns of OTUs revealed 81 OTUs with a higher frequency (p < 0.01) in one of the two different seasons (Figure 2). These comprised 54 OTUs during autumn season, with 7% of OTUs belonging to the phylum Endomyxa and 93% cercozoan OTUs. In spring, 27 cercozoan OTUs were detected to be particularly abundant. Taxonomic assignment of these OTUs identified OTU394 within the genus of *Rhogostoma* to be the most temporarily abundant OTU in autumn, followed by OTU627 assigned to the genus of *Thaumatomonas* and three endomyxan OTUs (OTU274, OTU230, OTU566) with >96% of their reads being found solely in autumn 2017 (Supplementary Figure S2, Supplementary Table S6). The endomyxan OTUs were root parasites (*Polymyxa betae*, OTU274; *Spongospora nasturtii,* OTU230) of the order Plasmodiophorida and a vampyrellid (OTU566), that were equally distributed across all canopy microhabitats and the ground in autumn. In spring, *Bodomorpha* sp. (OTU429), was temporarily highly abundant together with OTUs assigned to the genus *Thaumatomonas* (OTU472), two different *Euglypha* OTUs (OTU670, OTU675) and one *Paracercomonas* sp. (OTU735).

**Figure 2:**
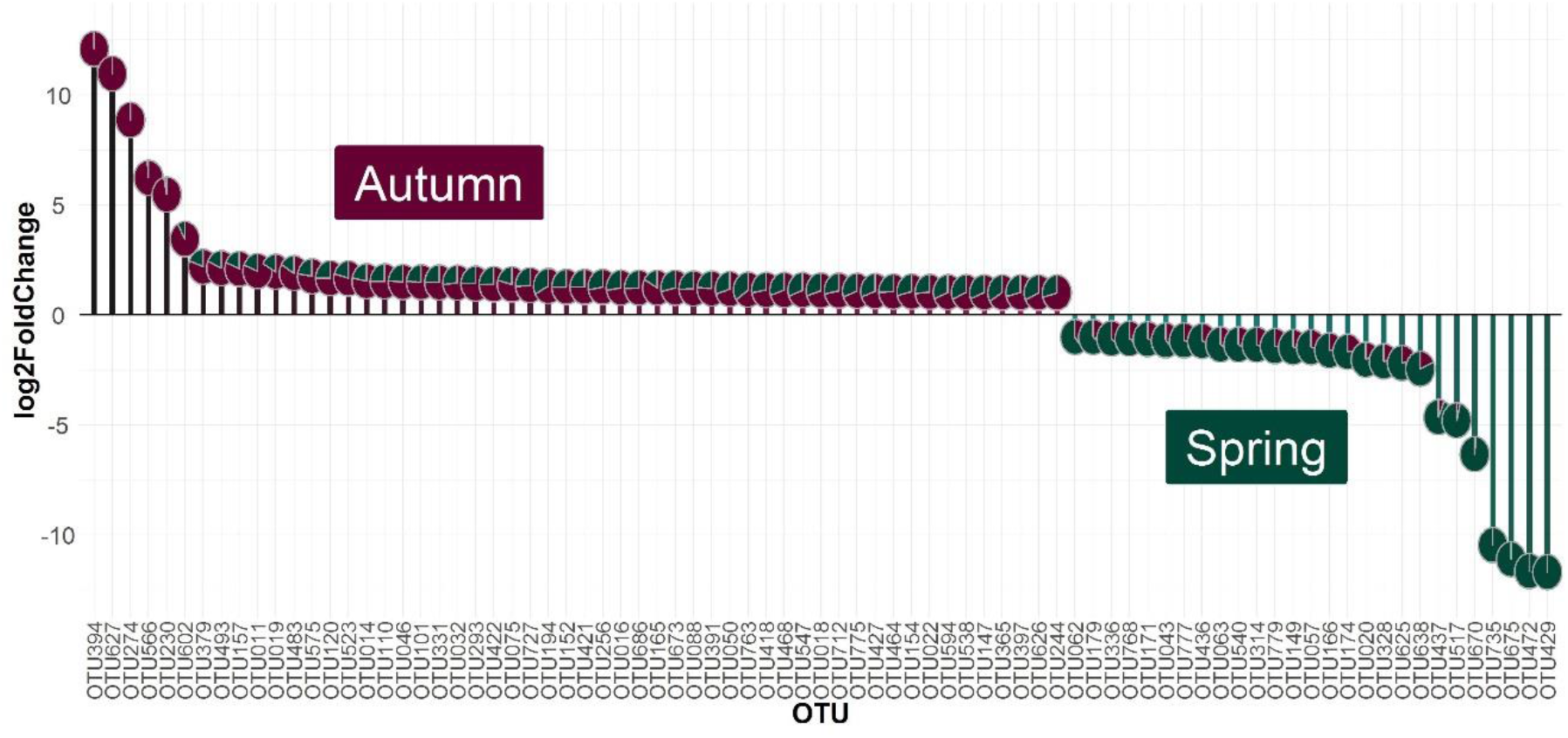
Analysis of season correlated OTUs. Investigation of autumn and spring communities revealed 54 and 27 OTUs with predominance in autumn and spring samplings, respectively (p < 0.01). Pie charts on the top of the bars represent the relative proportion of each OTU either in the autumn (purple) or spring (green) season.

Analysis of alpha diversity revealed similar patterns for every season (Supplementary Figure S3). However, OTU richness of fresh leaves showed much higher variation in spring as compared to autumn samples (ANOVA: F value = 5.98, p < 0.05); otherwise the general pattern was quite stable with fresh leaves, deadwood, arboreal soil and lichen having lower OTU richness as compared to bark and mosses *(Orthotrichum* sp., *Hypnum* sp.). On the ground, leaf litter appeared to have lower OTU richness then soil habitat (ANOVA: F value = 29.48, p < 0.001). In general, Simpson diversity, Shannon diversity, as well as species evenness showed almost the same pattern for both seasons (ANOVA; Simpson: F value = 3.55, p = 0.06; Shannon: F value = 0.28, p= 0.60; evenness: F value = 0.05, p = 0.82).

Non-metric multidimensional scaling of cercozoan and endomyxan beta diversity showed a clear separation between communities detected in the ground (litter and soil) and the canopy, plus a seasonal variability of these two strata (Figure 3, Supplementary Table S7). Most variation in protistan beta diversity within all four sampling periods was explained by microhabitat differences (perMANOVA: *R*^2^ 0.22, p < 0.01) and differences between canopy and ground (perMANOVA: *R*^2^ 0.17, p < 0.01). A very small, but significant proportion of beta diversity was explained by differences between the two seasons, spring and autumn (perMANOVA; canopy: *R*^2^ 0.02, p < 0.01; ground: *R*^2^ 0.05, p < 0.05). Tree species-specific differences between canopy communities were detected for *Quercus robur* (perMANOVA: *R*^2^ 0.04, p < 0.01) and *Tilia cordata* (perMANOVA: *R*^2^ 0.01, p < 0.01), although communities of fresh leaves were not influenced by the tree species (perMANOVA: *R*^2^ 0.11, p = 0.06), nor were the communities of leaf litter on the ground (perMANOVA: *R*^2^ 0.09 p = 0.10).

**Figure 3:**
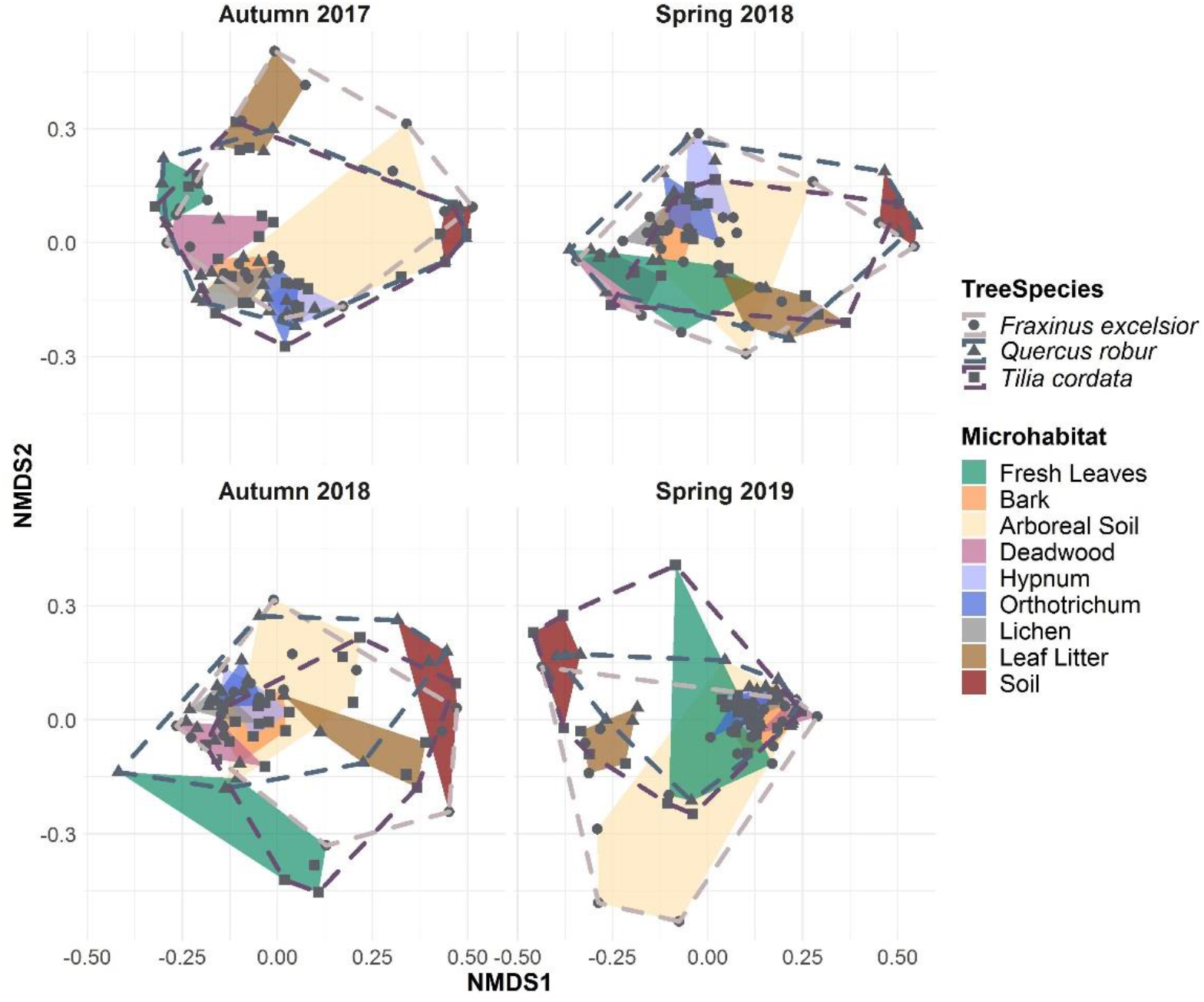
Non-metric multidimensional scaling (NMDS) of Bray-Curtis dissimilarities of cercozoan and endomyxan communities between microhabitats and tree species of each sampling period. Protistan communities showed a finer separation between canopy microhabitat communities during autumn, while leaf communities were more similar to other canopy microhabitat communities during spring (Stress values in Supplementary Table S9).

However, in autumn 2017 and spring 2018 cercozoan and endomyxan communities of leaf litter on the ground were more similar to the canopy communities than to the communities from the mineral soil directly underneath (Figure 3). Protistan communities detected on fresh leaves changed markedly between spring and autumn. In spring, communities detected on fresh leaves were still more similar to the other canopy microhabitats, but they became completely distinct in autumn (Supplementary Figure S4). Further, small seasonal differences in beta diversity for communities of bark and epiphytes with lichen and mosses *(Hypnum* sp. and *Orthotrichum* sp.) were detected (Supplementary Table S7). Communities of arboreal soil were highly variable in all four sampling periods, ranging from samples with high similarity to communities of the sampled epiphytes to communities closely resembling those of the mineral soil underneath the litter layer.

### Differentiation of foliar communities

Despite high differences in beta diversity between communities of all sampled microhabitats per sampling period (Figure 3), almost 98% of OTUs were shared between all sampling periods (Supplementary Figure S5). Accordingly, differences in community composition were almost entirely based on temporal and spatial changes in the relative abundance of species. Thus, principal coordinate analysis of all four sampling periods showed a high overlap of communities when taking all microhabitats into account (Figure 4 A), the first and second axis explained 25% and 11% of the variance, respectively. A separate analysis of fresh leaf samples only showed highly distinct autumn and spring communities (Figure 4 B; perMANOVA: *R*^2^ 0.15, p < 0.01). Both axes explained 38% of variance and not only separated the spring communities from autumn communities, but also showed no overlap between autumn communities of both years, suggesting a variable outcome after the recurrent community assembly over the seasons.

**Figure 4:**
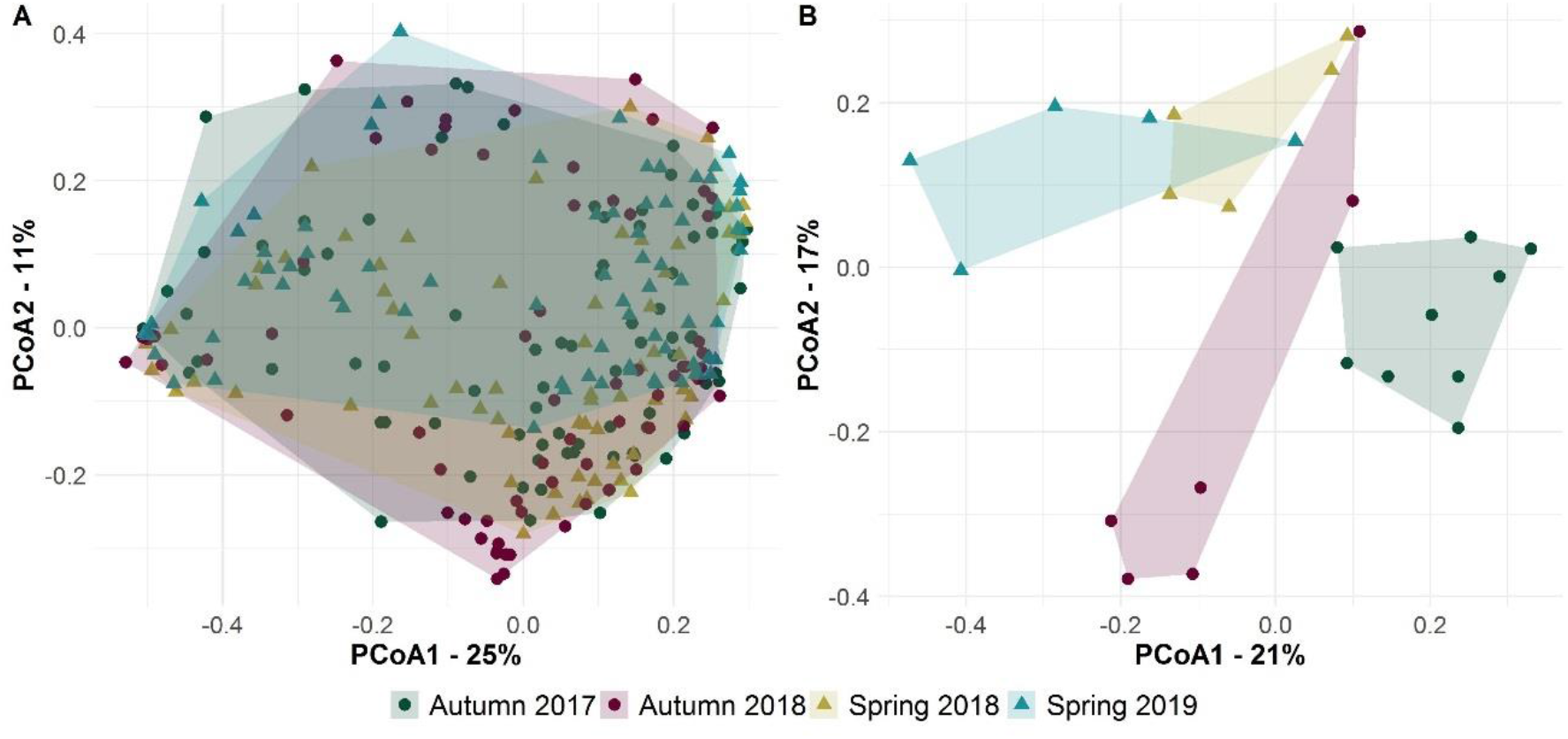
Principal Coordinates Analysis (PCoA) of cercozoan and endomyxan communities of all four sampling periods. Irrespective of the microhabitat identity, all sampling periods showed a comparable heterogeneity of detected communities (A). Cercozoan and endomyxan communities of fresh leaves where highly distinct between all four sampling periods, especially between the two seasons (B).

### Functional diversity

More than three-quarters of the canopy cercozoan and endomyxan reads were bacterivores (77 ± 9%), followed by omnivores (18 ± 7%), sequences of unknown function (4 ± 2%) and only very few eukaryvores (2 ± 1%) (Figure 5). Communities of ground microhabitats showed a relative smaller proportion of bacterivores (55 ± 11%; ANOVA: F = 31.09, p < 0.001) and more omnivores (26 ± 7%; ANOVA: F = 8.14, p < 0.01), aswell as a greater share of eukaryvores (5 ± 2%; ANOVA: F = 49.87, p < 0.001) compared to the canopy microhabitats. Plant parasites and parasites of other host organisms were only marginally present, on average <1%, except in autumn 2017, where soil communities contained 2.4% of reads derived from parasitic taxa. Most variation in protistan functional diversity was explained by differences between canopy and ground communities (perMANOVA: *R*^2^ 0.44, p < 0.01) and by microhabitat identity (perMANOVA: *R*^2^ 0.29, p < 0.01). However, functional group composition did not differ between seasons (perMANOVA; canopy: *R*^2^ 0.03, p = 0.37; ground: *R*^2^ 0.23, p = 0.24).

**Figure 5:**
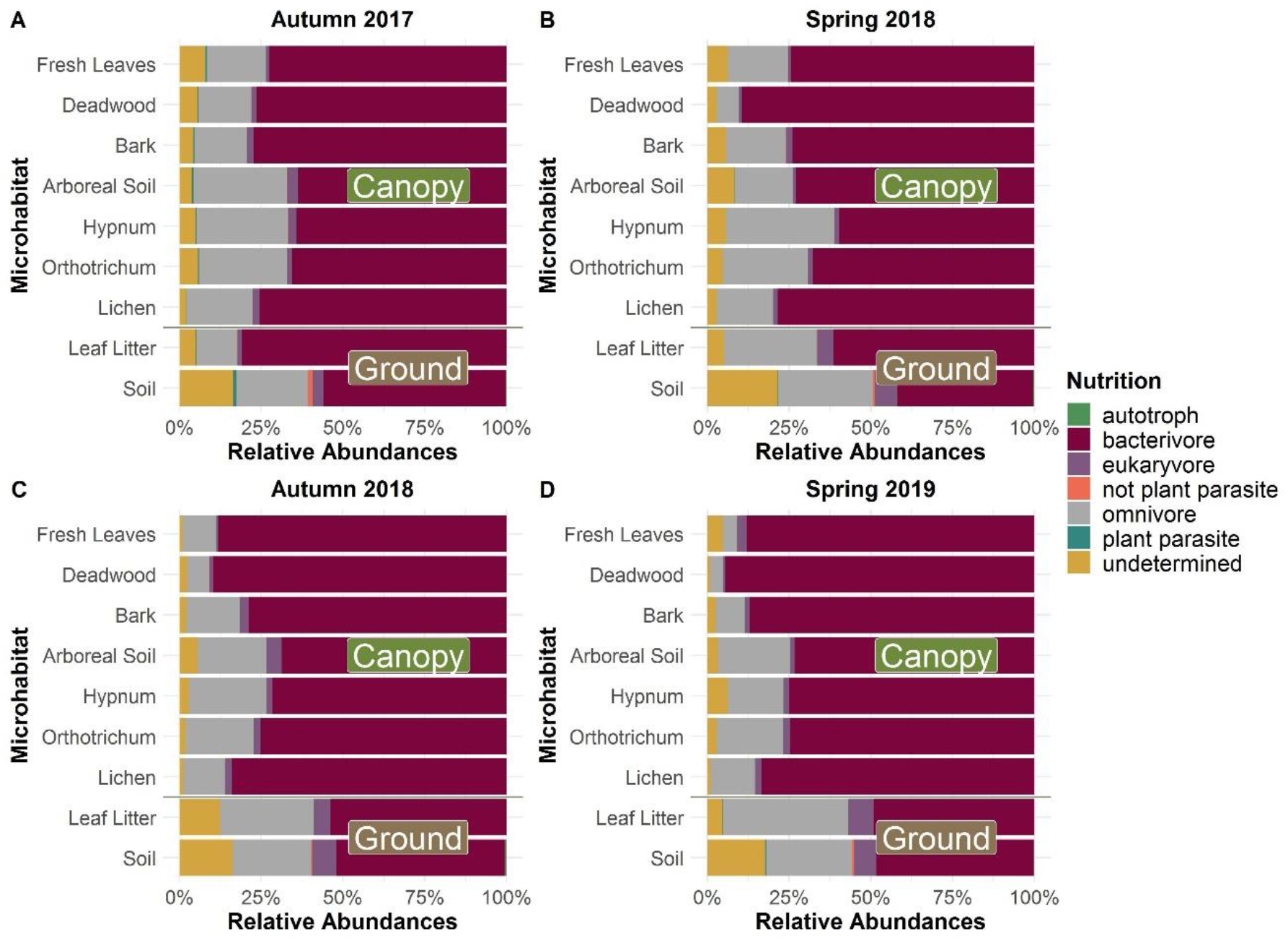
Relative read abundances of functional groups per sampled microhabitat and sampling period. Functional diversity of autumn (A,C) and spring samples (B,D) did not differ between seasons, but differences between sampled microhabitats and especially between the two strata (canopy and ground) were significant throughout all sampling periods. Bacterivores dominated, especially in tree canopies, whereas a higher proportion of omnivores and eukaryvores occurred on the ground.

## DISCUSSION

This study aimed to identify seasonal changes in the patterns of community composition of the diverse protistan phyla Cercozoa and Endomyxa, over two consecutive years. A total number of 783 OTUs were detected in the Leipzig floodplain forest, which is 43% of the cercozoan OTU richness that Fiore-Donno et al. (2020) retrieved with the same method from mineral soil of 150 different forest sites across Germany. The high sequencing depth, due to the taxon-specific primers (Fiore-Donno *et al.* 2018), enabled a direct comparison of protistan communities dwelling different microhabitats within the forest canopies. We showed that in principle all detected OTUs could be found everywhere in the floodplain forest, a pattern which was already described by Jauss et al. (2020a). However, patterns of cercozoan and endomyxan beta diversity in tree canopies were strikingly divergent from communities detected on the ground, showing that distinct species dominated the different communities. This was in particular true for the highly abundant glissomonad OTU001, with exceptionally higher relative abundance in canopies compared to the ground stratum. The clear differences between canopy and ground communities remained despite small, but significant seasonal changes.

### Seasonal variability of protists in tree canopies

Seasonality between spring and autumn explained 2% and 5% of the variation in beta diversity of canopy and ground communities, respectively (Figure 3, 4 B). About 10% of protistan OTUs were specifically associated with either spring or autumn season (Figure 2). For example, a *Rhogostoma* sp. (OTU394), belonging to omnivorous thecate amoebae in the Cryomonadida was temporally the most abundant taxon in autumn, while a bacterivorous *Bodomorpha* sp. from the order of Glissomonadida dominated in spring. Differences between spring and autumn communities were particularly evident on canopy leaves (Figure 4 B). In spring, beta diversity of the phyllosphere still showed some overlap with other canopy microhabitats (Figure 3). However, OTU richness showed very high variation and rarefaction curves of fresh canopy leaves did not reach a plateau (Supplementary Figure S1, S3), indicating high heterogeneity during community assembly shortly after leaf emergence in spring, while the distinct separation of beta diversity in autumn shows that specific leaf surface communities had established (Figure 3 B, 4 B). However, beta diversity of fresh leaves communities showed no overlap between both autumn samplings (Figure 4 B), indicating variable outcomes of community assembly driven by seasonal factors. October 2017 was an exceptionally warm and wet month, while October 2018 and the prior season was too warm and exceptionally dry (DWD; 2017, 2018). Nevertheless, autumn samples explained much more variation in beta diversity than spring samples (Supplementary Figure S4). Especially in 2017, ordination placed beta diversity of leaf litter communities on the ground between soil and foliar communities in the phyllosphere (Figure 3), suggesting that leaf litter still carries a signature of the preceding foliar community (Jauss *et al.* 2020a). Our environmental sequencing method, based on ribosomal DNA, did not allow to distinguish between active protists and their resting or dispersal stages, but instead must be considered as an integrative long-term measure of taxa that replicated well and formed resting stages in respective microhabitats. The clear differences in beta diversity between microhabitats indicate that well-adapted taxa accumulated and dominated over those that arrived as resting stages by passive dispersal. This leads to differences in traits, which only can be inferred from related taxa (Dumack *et al.* 2020) as almost 80% of the OTUs showed a similarity of less than 97% to any sequence in the reference database, confirming the existence of a substantial undescribed taxonomic diversity within this dominant phylum of microbial eukaryotes in terrestrial ecosystems (Singer *et al.* 2021).

### Protistan diversity and functional traits

The majority of the 783 OTUs could be assigned to the phylum Cercozoa (97%), the remaining to Endomyxa (3%) and to the incertae sedis Novel clade 10 (Tremulida <1%) (Supplementary Figure S6). With 753 OTUs cercozoan diversity was in line with previous studies, which established Sarcomonadea (Glissomonadida and Cercomonadida) as the dominant class in terrestrial habitats (Geisen *et al.* 2015; Ploch *et al.* 2016; Fiore-Donno *et al.* 2018). Especially the small and bacterivorous flagellates in the order Glissomonadida dominated throughout all canopy microhabitats during all four sampling periods (Figure 1, 5, Supplementary Figure S7). The Sarcomonadea were followed by mainly omnivorous testate amoebae in the orders Euglyphida and Cryomonadida. These omnivores can feed on both, bacteria and small eukaryotes, such as yeasts, algae and other protists (Dumack *et al.* 2020). While bacteria appeared as an essential food source in tree canopies, cercozoan communities of litter and mineral soil on the ground were characterized by a higher proportion of eukaryvores, which was mostly related to higher relative read numbers of vampyrellid amoebae that feed on a wide range of soil eukaryotes, including fungal mycelia and spores, algae, as well as nematodes (Anderson and Patrick 1980; Surek and Melkonian 1980; Hess, Sausen and Melkonian 2012). Our findings reflect the results of Fiore-Donno et al. (2020), who found a high proportion of vampyrellids, but almost no other Endomyxa in mineral soil samples of diverse forests in different regions in Germany. In addition, reads derived from taxa of so far undetermined feeding mode were enriched in litter and soil compared to canopy samples (Figure 5), indicating a more complex structure of microeukaryote food webs on the ground than in the physically harsh environment of the tree crown.

Most variation in cercozoan and endomyxan functional diversity was explained by microhabitat differences and the differences between canopy and ground communities, whereas seasonality with respect to the investigated functional traits was not observed. However, seasonal differences could be revealed when taking taxonomically assigned relative read abundances into account (Supplementary Figure S7). One explanation for this pattern is that the abundance of less dominant orders was more variable between the sampled microhabitats and seasons. Because the functional traits (especially feeding traits) are still understudied, a measurable proportion of traits could not be assigned to the detected taxa (Canopy: 4 ± 2%, Ground: 12 ± 6%).

### Forest canopies as a reservoir for potential plant pathogens

Spatial distribution of endomyxan plant parasites is patchy and increasing evidence hints to the habitat type as primary explanatory force. Khanipour Roshan et al. (2021) found them to be almost entirely missing in the surface of sandy dunes. Further, Fiore-Donno, Richter-Heitmann and Bonkowski (2020) reported an almost complete absence of endomyxan plant parasites in forest soils. We, however, found two temporally abundant OTUs among autumn communities which could be assigned to the root pathogenic species of *Polymyxa betae* and *Spongospora nasturtii* (Phytomyxea: Plasmodiophorida) (Figure 2). Whereas, *S. nasturtii* is an obligate biotrophic root pathogen of watercress (*Nasturtium officinale)* (Down, Grenville and Clarkson 2002), a common herb of river banks in the floodplain forest. *P. betae* is an obligate root parasite in beet roots (Tamada and Asher 2016), and although its potential host range also includes Chenopodiaceae, Caryophyllaceae and Papaveraceae, (Barr and Asher 1992; Neuhauser *et al.* 2014), none of these host plants were detected in the sampled forest. The ubiquitous distribution of these two plasmodiophorids among protists of tree crowns, litter and soil in autumn samples (Supplementary Figure S2) reflects the complex life cycle of these plant pathogens with distribution via sporangia in autumn (Barr and Asher 1996). The high potential of wind dispersal of protistan propagules, was recently emphasized by Jauss et al. (2020b) and together with our results it appears that tree canopies play potential role as reservoirs for plant pathogenic microbial propagules. Or the other way around: tree canopies play a potential role as physical filters that may partly prevent the further spread of these plant pathogens.

## Conclusion

Investigation into two important protistan lineages, Cercozoa and Endomyxa, over a period of two years revealed strong differences in community composition between canopy and soil microhabitats, and a small, but significant fraction of recurrent seasonal variability of these communities. We observed lower beta diversity between canopy communities in spring compared to autumn. Especially foliar communities changed during the aging of leaves from spring to autumn, indicating an interannual community assembly. One particular glissomonadid OTU appeared to be a canopy specialist, while high read numbers of root parasitic phytomyxean OTUs in tree canopies during autumn demonstrate a potential role of the canopy surface as an important reservoir for wind-dispersed propagules of microbial eukaryotes. Occasionally leaf litter communities showed more similarity to foliar canopy communities than to those of the soil directly underneath. Thus, after litter fall, the preceding seasonal community assembly in the canopy contributes to spatial differences of protistan communities on the ground, but the latter become enriched in omnivores and eukaryvores relative to the predominantly bacterivorous canopy inhabitants. The described diversity of Cercozoa and Endomyxa in this study is just one striking example of dozens of microbial eukaryote phyla whose canopy inhabitants still await discovery.

## Acknowledgements

The authors would like to thank Rolf Engelmann for his assistance with the field work by operating the canopy crane, as well as the Leipzig Canopy Crane Platform of the German Centre for Integrative Biodiversity Research (iDiv) for providing the site access and allowing us to sample the trees from their field trial.

## Data Accessibility

Raw sequence data have been submitted to the European Nucleotide Archive (ENA) database under the Bioproject number PRJEB37525, with run accession numbers ERR3994029, ERR4913261 and ERR4911998.

## Author contributions

MB and MS designed the study. SW, R-TJ, SS, RW and KF conceived and conducted the sampling and DNA extraction. AMF-D contributed the primers. KD helped in laboratory work. SW and KF conducted the PCRs. R-TJ assisted in bioinformatics, provided the original R script to detect temporally abundant OTUs. SW performed the bioinformatic and statistical analyses and drafted the manuscript. All authors contributed to and approved the final version.

