## Supplementary for "On the phenology of protists: Recurrent patterns reveal seasonal variation of protistan (Rhizaria: Cercozoa, Endomyxa) communities in tree canopies"

### *Supplementary Figures*

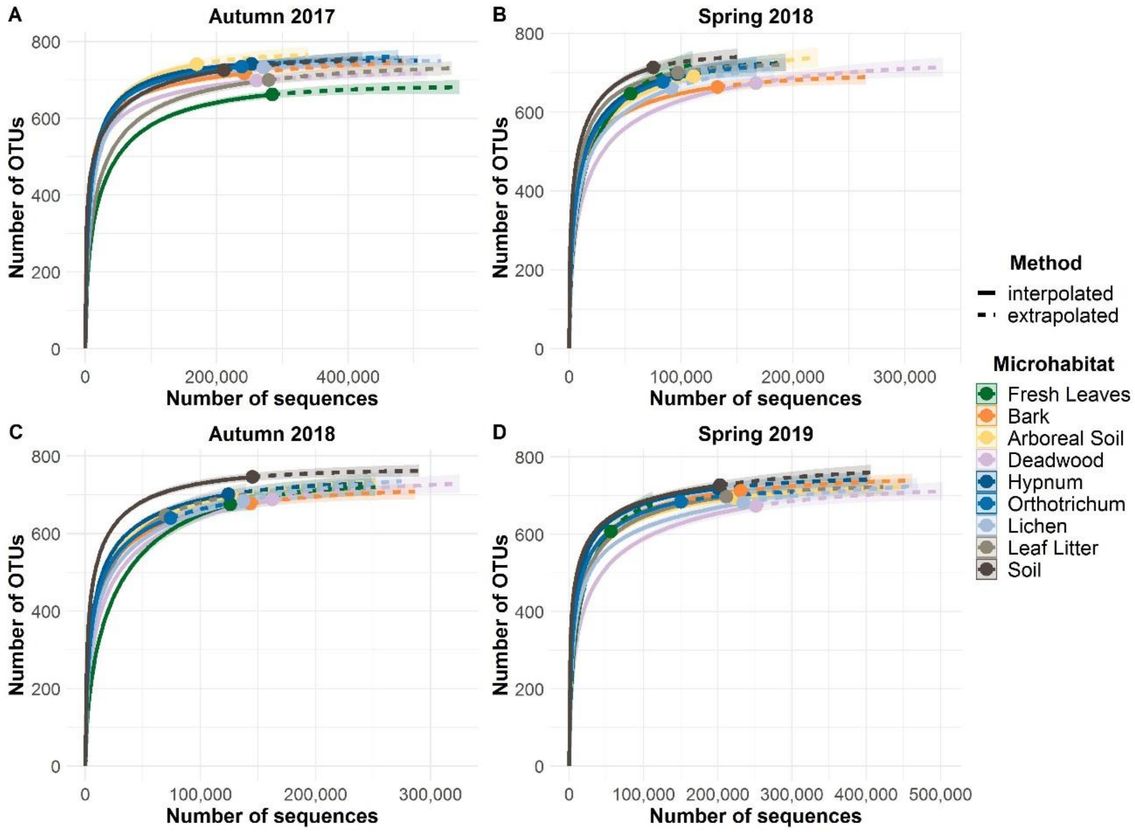


**Supplementary Figure S1. Rarefaction curves of each sampling period.** Solid lines give the interpolated number of OTUs from subsampled sequences, dashed lines represent extrapolated number of OTUs with increasing number of sequences. Shaded areas give the 97% confidence intervals.

**On the phenology of protists: Recurrent patterns reveal seasonal variation of protistan (Rhizaria: Cercozoa, Endomyxa) communities in tree canopies**

Susanne Walden, Robin-Tobias Jauss, Kai Feng, Anna Maria Fiore-Donno, Kenneth Dumack, Stefan Schaffer, Ronny Wolf, Martin Schlegel, Michael Bonkowski

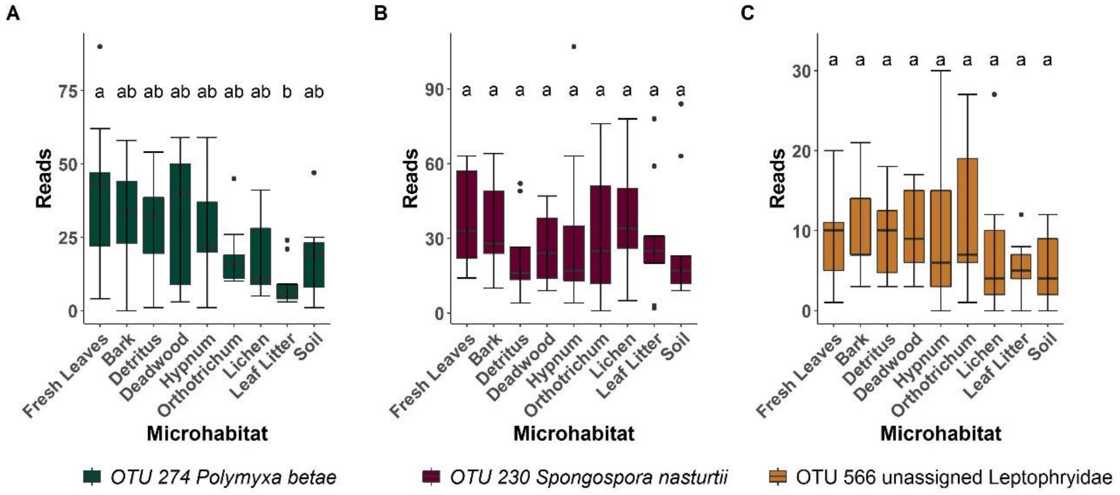


**Supplementary Figure S2. Boxplot of most differential abundant endomyxan OTUs in autumn 2017.** Boxplots describe the absolute contribution of reads from three highly differential abundant endomyxan OTUs detected in all microhabitats in autumn 2017 - outliers are given by dots. Letters correspond to Tukey's Honest Difference post hoc test, with microhabitats not sharing any letter having significantly different means.

**On the phenology of protists: Recurrent patterns reveal seasonal variation of protistan (Rhizaria: Cercozoa, Endomyxa) communities in tree canopies**

Susanne Walden, Robin-Tobias Jauss, Kai Feng, Anna Maria Fiore-Donno, Kenneth Dumack, Stefan Schaffer, Ronny Wolf, Martin Schlegel, Michael Bonkowski

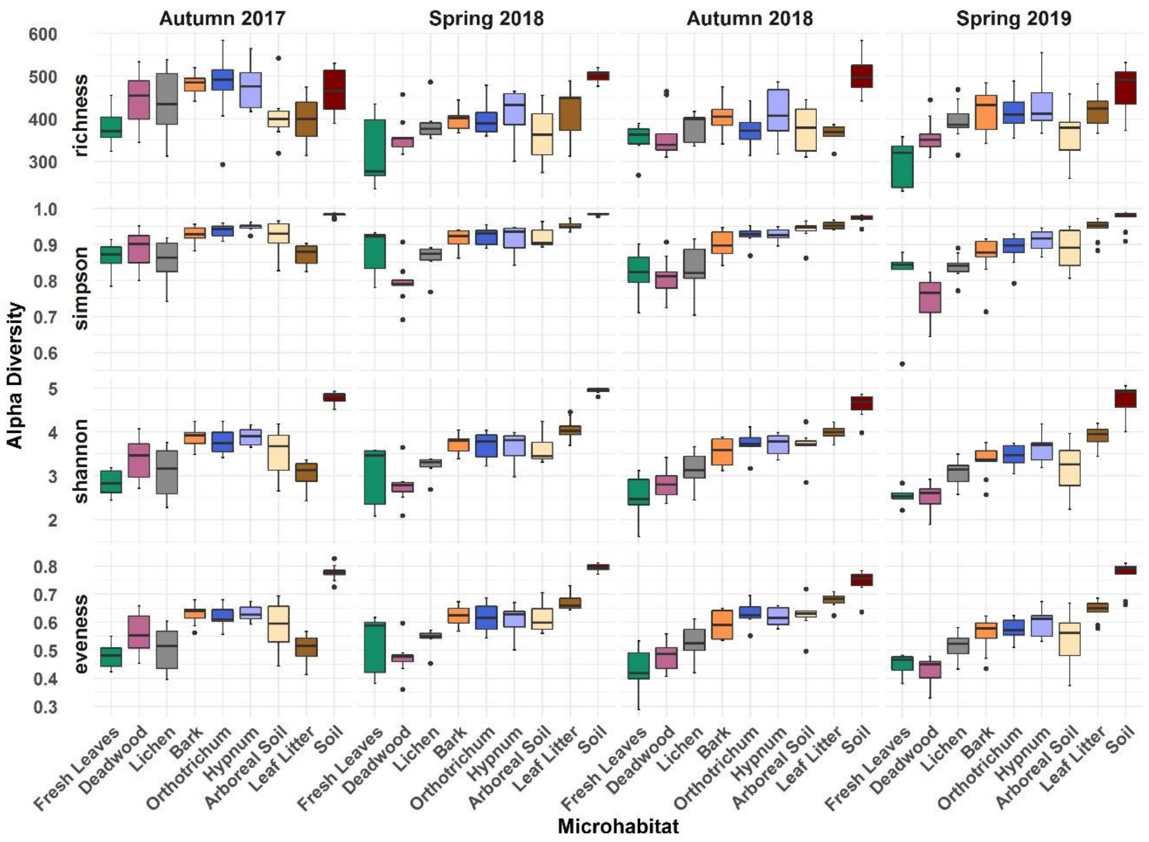


**Supplementary Figure S3. Alpha diversity of microhabitats for cercozoan and endomyxan communities of four different sampling periods.** Boxplots describe the species richness, Simpson Index, Shannon Index and evenness (in descending order) of the microhabitats grouped by sampling period, outliers are given by dots.

**On the phenology of protists: Recurrent patterns reveal seasonal variation of protistan (Rhizaria: Cercozoa, Endomyxa) communities in tree canopies**

Susanne Walden, Robin-Tobias Jauss, Kai Feng, Anna Maria Fiore-Donno, Kenneth Dumack, Stefan Schaffer, Ronny Wolf, Martin Schlegel, Michael Bonkowski

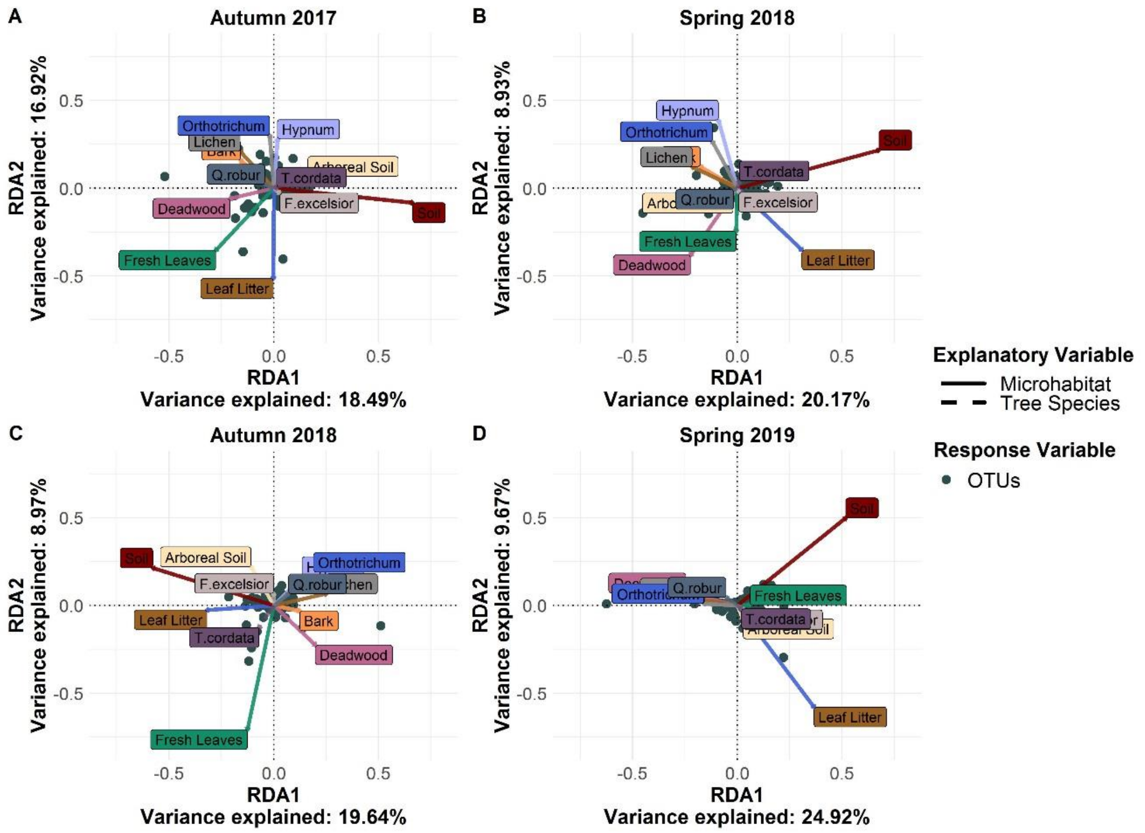


**Supplementary Figure S4. Redundancy analysis (db-RDA) of cercozoan and endomyxan OTUs of four sampling periods.** Environmental factors of microhabitat identity and tree species were included in the analysis. Dots represent OTUs. The percentages of variability explained by each axis (RDA1 and RDA2) are given in the labels. RDA ordination resulted in more distinct fresh leaves communities in autumn.

**On the phenology of protists: Recurrent patterns reveal seasonal variation of protistan (Rhizaria: Cercozoa, Endomyxa) communities in tree canopies**

Susanne Walden, Robin-Tobias Jauss, Kai Feng, Anna Maria Fiore-Donno, Kenneth Dumack, Stefan Schaffer, Ronny Wolf, Martin Schlegel, Michael Bonkowski

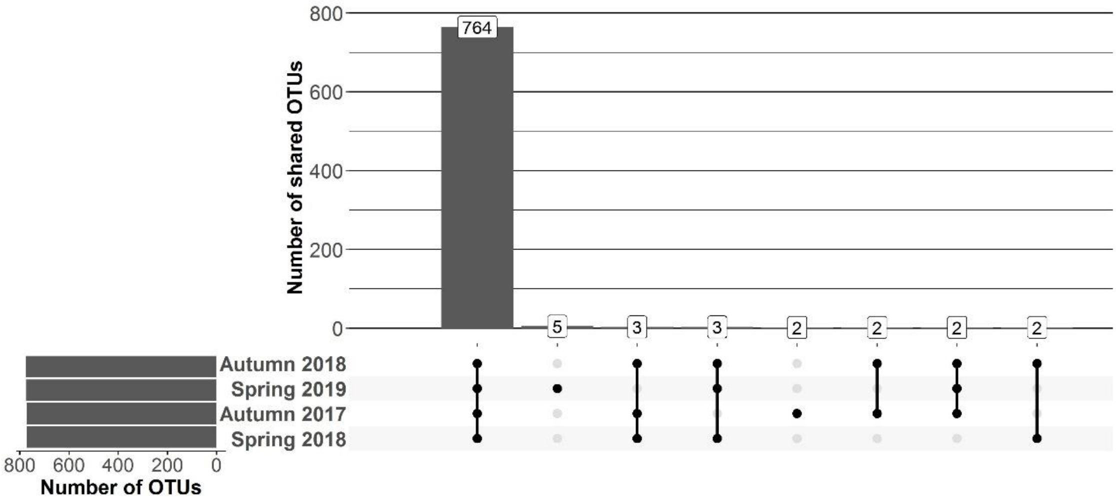


**Supplementary Figure S5. Shared OTUs of cercozoans and endomyxans between sampling periods.** Top bar charts represent the sum of the number of shared OTUs resulting from the combination of sampling periods in the matrix below. Left bar charts show the number of OTUs detected in every sampling period in descending order. Of all OTUs 98% (764) are shared between all four samplings.

**On the phenology of protists: Recurrent patterns reveal seasonal variation of protistan (Rhizaria: Cercozoa, Endomyxa) communities in tree canopies**

Susanne Walden, Robin-Tobias Jauss, Kai Feng, Anna Maria Fiore-Donno, Kenneth Dumack, Stefan Schaffer, Ronny Wolf, Martin Schlegel, Michael Bonkowski

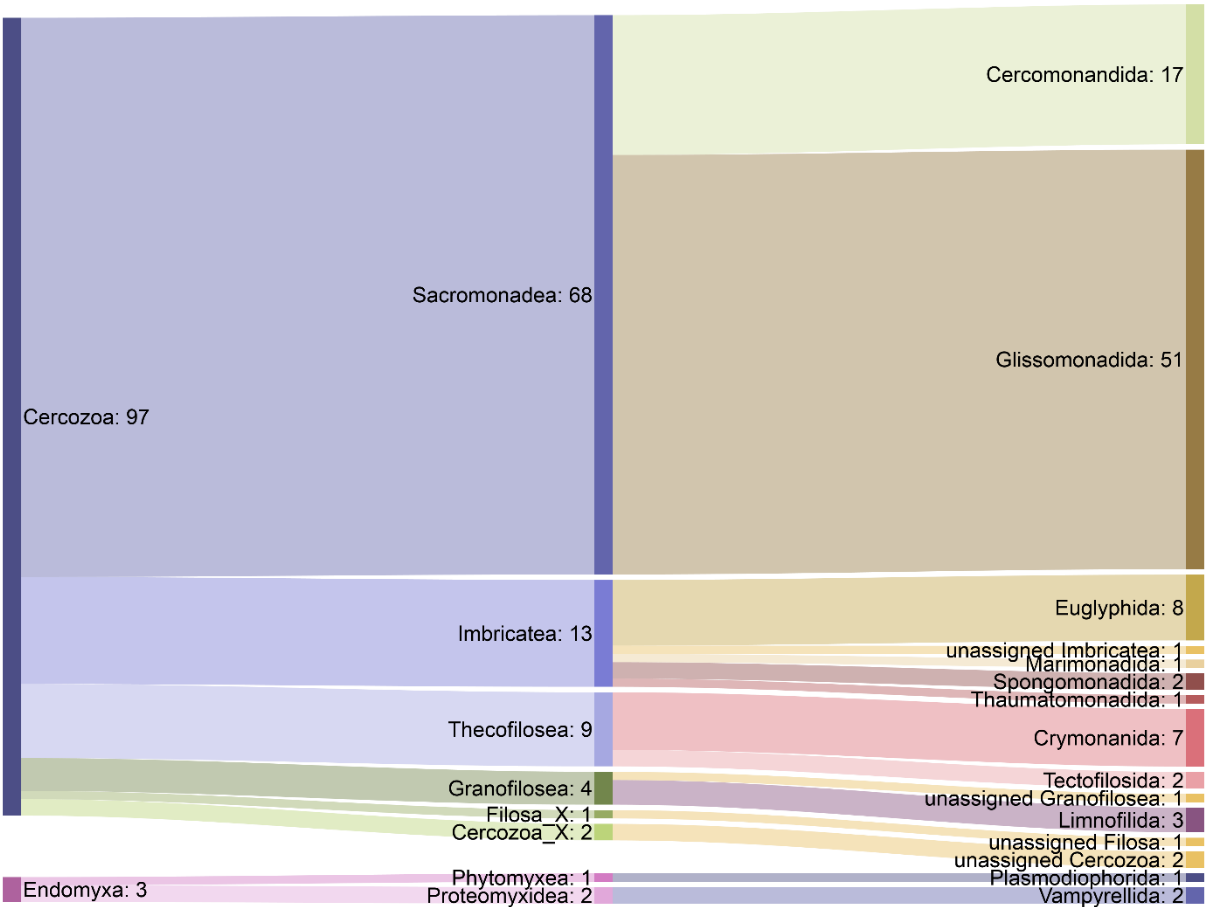


**Supplementary Figure S6. Sankey diagram showing the summarized relative abundances of the cercozoan and endomyxan OTUs to the taxonomic diversity.** Taxonomical assignment is based on the best hit by BLAST. From left to right, names refer to phylum (Cercozoa, Endomyxa), class and orders. Taxa represented by <1% are not depicted for the sake of clarity.

**On the phenology of protists: Recurrent patterns reveal seasonal variation of protistan (Rhizaria: Cercozoa, Endomyxa) communities in tree canopies**

Susanne Walden, Robin-Tobias Jauss, Kai Feng, Anna Maria Fiore-Donno, Kenneth Dumack, Stefan Schaffer, Ronny Wolf, Martin Schlegel, Michael Bonkowski

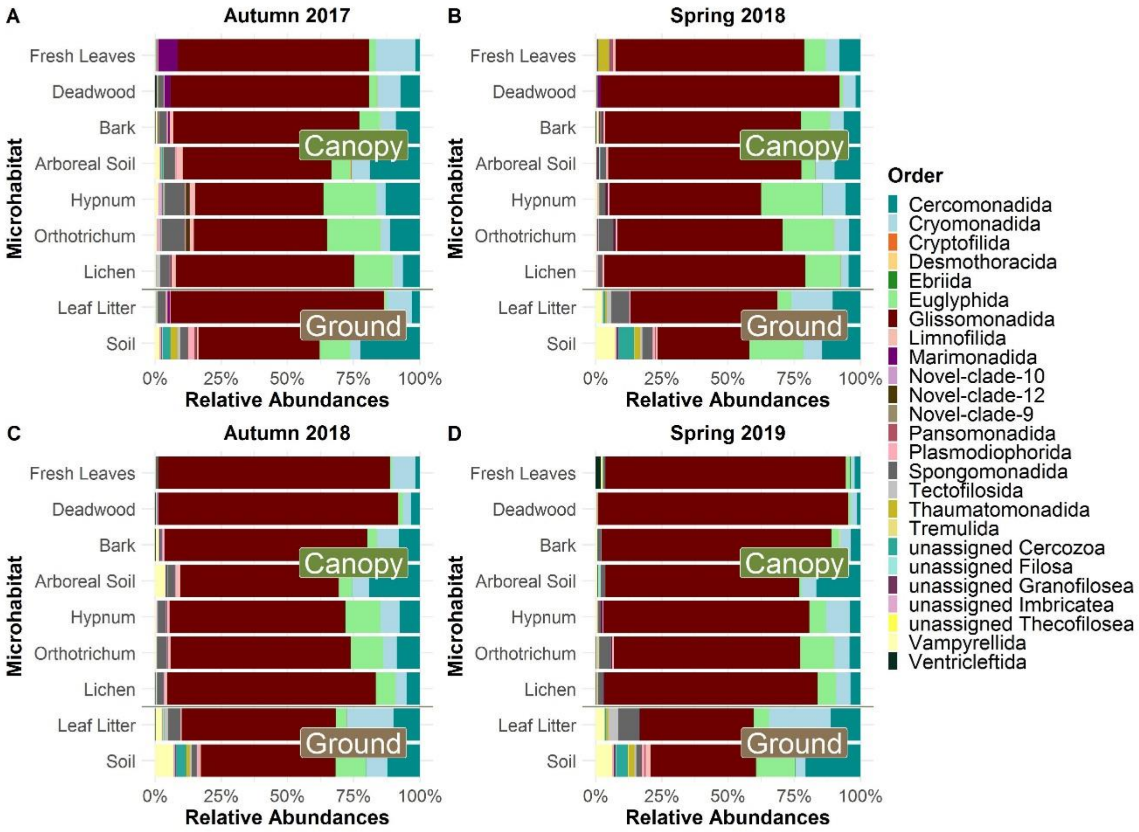


**Supplementary Figure S7. Relative read abundances of cercozoan and endomyxan orders per sampled microhabitat and sampling period.** Number of reads of the order Glissomonadida where most abundant throughout all sampling periods and microhabitats. Relative read abundances of predatory Vampyrellida occurred with up to 7% in sampled mineral soils.
